# Potential Functions of Histone H3.3K56 Acetylation in Mammals

**DOI:** 10.1101/2020.03.16.994616

**Authors:** Lei Fang, Danqi Chen, Jingzi Zhang, Hongjie Li, Beatrix Bradford, Chunyuan Jin

## Abstract

H3K56 acetylation (H3K56Ac) was first identified in yeast and has recently been reported to play important roles in maintaining genomic stability, chromatin assembly, DNA replication, cell cycle progression and DNA repair. Although H3.1K56Ac has been relatively well studied, the function of H3.3K56Ac remains mostly unknown in mammals. In this study, we used H3.3K56Q and H3.3K56R mutants to study the possible function of H3.3K56 acetylation. The K-to-Q substitution mimics a constitutively acetylated lysine, while the K-to-R replacement mimics a constitutively unmodified lysine. We report that cell lines harboring mutation of H3.3K56R exhibit dramatic morphology changes and cell death. Using Tet-Off inducible system, we show an increased population of polyploid/aneuploid cells and a decreased cell viability in H3.3K56R mutant cells. In consistence with these results, H3.3K56R mutant compromised H3.3 incorporation into several pericentric and centric heterochromatin regions we tested. Moreover, mass spectrometry analysis coupled with label free quantification reveal that biological processes regulated by the H3.3-associating proteins, whose interaction with H3.3 is markedly increased by H3.3K56Q mutation but decreased by H3.3K56Q mutation, include sister chromatid cohesion, mitotic nuclear division, and mitotic nuclear envelope disassembly. These results suggest that H3.3K56 acetylation is crucial for chromosome segregation and cell division in mammals.

## Introduction

A Nucleosome is the basic unit of chromatin in eukaryotes, consisting of a histone octamer wrapped by 147 bp DNA [1, 2]. A histone octamer contains two copies of four different types of core histones H2A, H2B, H3, and H4 [3, 4]. In addition to canonical histones, several histone variants such as H2A.Z, H2A.X, and H3.3 have also been identified, which play pivotal roles in modulating chromatin dynamics and in the activities of the underlying DNA [5–8]. Among these histone variants, H3.3 is well-known for its functions in regulating gene expression, DNA replication, and DNA repair, as well as its roles in cell cycle control, cell differentiation, development, and cancer [7, 9–13]. H3.3 was initially identified localized at active genomic loci such as gene bodies of active genes, active promoters and regulatory regions [8, 10, 14–16]. A number of studies later demonstrated that H3.3 is also deposited at heterochromatin regions such as pericentric heterochromatin and telomere [11, 17–21]. The function of H3.3 at these regions as well as the post-translational modification(s) important for the H3.3 incorporation have remained elusive.

All histones contain a globular core domain and more relaxed N-terminal and C-terminal tails. Lysine residues on both globular domain and tails of histones can be modified by post-translational modifications (PTMs) such as acetylation, phosphorylation, ubiquitination and methylation [2, 22–26]. These histone modifications are related to the different states of chromatin and regulate distinct histone-involved processes, including DNA replication, gene transcription and DNA repair [3, 5, 7, 13]. Histone modifications also affect chromatin assembly in various ways, including the regulation of histone folding and processing, histone nuclear import, and the interaction between histones and histone chaperones [25, 27–29].

Histone H3 lysine 56 (H3K56) is a core domain residue localizing at both the entry and exit point of a nucleosome [30, 31]. H3K56 acetylation has been identified in yeast, *Drosophila* and mammals, where it has been shown to play important role in maintaining genomic stability, DNA replication, cell cycle progression and DNA repair [32–42]. H3K56Ac is very abundant in yeast, marking ~30% of total histone H3. However, in mammalian cells its abundance is much lower as it marks less than 1% of total H3 [34]. H3K56Ac is tightly regulated by cell cycle and DNA damage-induced checkpoints [30, 32]. In budding yeast, H3K56 has been reported to be acetylated predominantly during the S phase, but deacetylated rapidly when cells enter the G2/M phase of the cell cycle [32, 43]. Mammalian H3K56 acetylation requires the histone chaperone Asf1 and occurs mainly at the S phase in unstressed cells [37, 42]. Dysregulation of histone H3K56 acetylation leads to increased sensitivity to DNA damage agents and elevated genome instability [37, 44, 45]. Both hyper- and hypo-acetylation of H3K56 result in defects in sister chromatid cohesion, recombination and ribosomal DNA (rDNA) amplification [27, 33]. H3K56Ac makes the nucleosome termini more flexible and facilitates nucleosome disassembly [41, 46, 47]. H3K56Ac also alters the substrate specificity of SWR-C, leading to either H2A.Z or H2A being exchanged from nucleosomes [48]. These findings strongly support the important role of H3K56 acetylation in regulating cell cycle progression, DNA damage response, chromatin remodeling, nucleosome assembly following DNA replication and DNA repair. However, all these reports focused on K56Ac of H3, either on H3.1 or without defining which isoform of H3. Studies on K56Ac of the variant H3.3 in mammals in particular are absent and remain to be explored.

In this study, to address the function of H3.3K56 acetylation, we made use of H3.3K56Q and H3.3K56R mutants, in which the lysine at position 56 was converted to glutamine (Q) and arginine (R) respectively. The K-to-Q substitution mimics lysine acetylation, whereas the K-to-R replacement prevents lysine acetylation. We studied the effects of H3.3 acetylation on cell growth, cell cycle progress, DNA damage response, nucleosome assembly, and protein association in human cells. The results suggest that H3.3K56 acetylation is crucial for chromosome segregation and cell division as well as for cell cycle progress and nucleosome assembly in mammals, possibly through changing the spectrum of H3.3-associating proteins. These findings add insights into the function of H3.3K56 acetylation in mammals.

## Materials and methods

### Plasmids and cell culture

A cDNA fragment of H3.3 was amplified by PCR using pcDNA3.1-FLAG-H3.3 as a template. The primers used were: Forward: 5’-CAGATCTATGGCTCGTACGAAGCAAAC-3’ and Reverse: 5’-ATCTAGACTAGGCGTAGTCGGGCACG-3’. The fragment was cloned into the Bgl II/Xba I sites of puHD-FLAG to generate puHD-FLAG-H3.3WT construct.

The plasmid pUHD-FLAG-H3.3K56R and pUHD-FLAG-H3.3K56Q were constructed using a Quick Change II Site-Directed Mutagenesis Kit (Agilent, CA, United States), according to the manufacturer’s instructions, with the primers H3.3K56R-Forward: 5’-GAGATTCGTCGTTATCAG***CG6***TCGACCGAGCTGCTCAT-3’ and H3.3K56R-Reverse: 5’-ATGAGCAGCTCGGTCGA***CC6***UTGATAACGACGAATCTC-3’; H3.3K56Q-Forward: 5’-GAGAGATTCGTCGTTATCAG***C46***TCGACCGAGCTG-3’ and H3.3K56Q-Reverse: 5’-CAGCTCGGTCGA***CTG***CTGATAACGACGAATCTCTC-3’; the bold italic letters represent the mutant nucleotide. The presence of the desired mutation was confirmed using nucleotide sequencing (Invitrogen, CA, United States).

Similarly, pUHD-FLAG-H3.1K56R and pUHD-FLAG-H3.1K56Q were constructed using H3.1 cDNA as template with the primers H3.1K56R-Forward: 5’-CGTCGCTACCAG***A6V***ΓCCACCGAGCTG −3’ and H3.1K56R-Reverse: 5’-CAGCTCGGTGGA***CCT***CTGGTAGCGACG −3’; H3.1K56Q-Forward: 5’-CCGTCGCTACCAG***C46***TCCACCGAGCT −3’ and H3.1K56Q-Reverse: 5’-AGCTCGGTGGA***CT6***UTGGTAGCGACGG −3’; the italic bold letters represent the mutant nucleotide.

The plasmids were separately transfected into UTA6 human osteosarcoma cells, which contain tetracycline transactivator tTA, with Lipofectamine 2000 (Invitrogen, CA, United States), and selected with zeocin (60 μg/ml) and tetracycline (1 μg/ml) to generate inducible UTA6-FLAG-H3.3WT/K56Q/K56R stable cell lines. These UTA6-FLAG-H3.3 stable cell lines were cultured in Dulbecco’s modified Eagle’s medium (DMEM) supplemented with 10% fetal bovine serum, 100 U/ml penicillin, and 100 μg/ml streptomycin. Cells were incubated at 37°C in a humidified atmosphere of 5% carbon dioxide.

### Antibodies

The antibodies were purchased as following: anti-Histone H3 (ab1791), anti-gamma H2A.X (ab11174), and Anti-Actin (ab8226) were from Abcam; anti-FLAG (F3165) was from Sigma.

### FACS analysis of mutant cells

The UTA6-FLAG-H3.3WT/K56Q/K56R stable cell lines were grown in zeocin-/tetracycline-DMEM to induce the expression of H3.3K56 mutants for 3, 4 or 5 days, and then collected for FACS analysis. Briefly, ~3×10^6^ cells were fixed in ice cold 70% ethanol and stored at −20°C for later analysis. The fixed cells were centrifuged at 1,000 rpm for 10 min and washed with cold PBS twice. Propidium iodide (PI) staining solution (50 μg/ml final concentration; Acros Organics) was added to the cells and incubate for 30 min at room temperature in the dark. Next, 100,000 cells were analyzed using Epics SL-MCL (BECKMAN COULTER) equipped with a suitable laser and EXPO32 ADC Analysis software.

### Cell viability assay

Cell viability assay was performed using MTT method to test if the cell proliferation has changed in mutant cells. 8×10^3^ cells of each cell line were seeded in one well of 96-well plate, and grown for 0, 1, 2, 3, 4, 5 or 6 days before measuring their cell viability. Cell Titer 96®Aqueous One Solution Reagent (MTS assay, Promega, Catalog G3582) was added to each well according to the manufacturer’s instructions. After 3-hr incubation, cell viability was determined by measuring the absorbance at 490 nm using a 550 BioRad platereader (Bio-Rad, Hertfordshire, UK). At least triplicates were done for each condition to get statistical significance. Day 0 cells were used as control for normalization.

### Treatment of DNA damage reagents

H3K56Ac was reported to be essential in DNA damage response. To investigate the role of H3.3K56Ac in DNA damage, the mutant cells were treated with different dose of capmtothecin (CPT, Sigma, St. Louis, MO) and methyl methanesulfonate (MMS, Sigma, St. Louis, MO) and their cell viabilities were measured using MTT method. Non-treated cells were used as control for normalization. Cells were plated on 96-well plates with normal medium at a density of 8×10^3^ cells/well. After overnight incubation, cells were treated with CPT or MMS at the indicated dosages. After 24 h (MMS) or 48 h (CPT) incubation, cell viability was determined by the MTS assay mentioned above.

### Cell synchronization and release from thymidine block

To clarify the function of H3.3K56Ac in modulating cell cycle progression, we synchronized H3.3WT/ K56Q/ K56R cells at G1/S using double thymidine block, and then release the cells to different cell phases to monitor if there are any cell cycle progression defects. Briefly, cells were seeded in 30% confluence and grown for 24h before adding 2mM thymidine (final concentration) to the medium for the first block. After 24h, the medium was discarded and the cells were washed with PBS for 3 times, before fresh medium was added to release at 12h.Then 2mM thymidine was added for another 16h for the second block at G1/S border. The cells were then released by replacing with fresh medium for 2, 5, 8, 12h to enter different cell phases, and DNA contents were determined by FCAS analysis.

### Nucleosome preparation and ChIP-qPCR

Mono- and di-nucleosomes were isolated by Micrococcal Nuclease (MNase) digestion and sucrose gradient purification. Chromatin immunoprecipitation (ChIP) analysis was performed as described previously [49]. Briefly, 50 μg chromatin was used for ChIP with anti-FLAG M2 agarose beads (A2220, Sigma, Sigma, St. Louis, MO), and the obtained ChIP-DNA was subject to qPCR using primers representing different regions of genome, including enhancer/promoters, transcriptional termination site, rDNA regions, pericentric and centric heterochromatin regions.

The following primers were used:

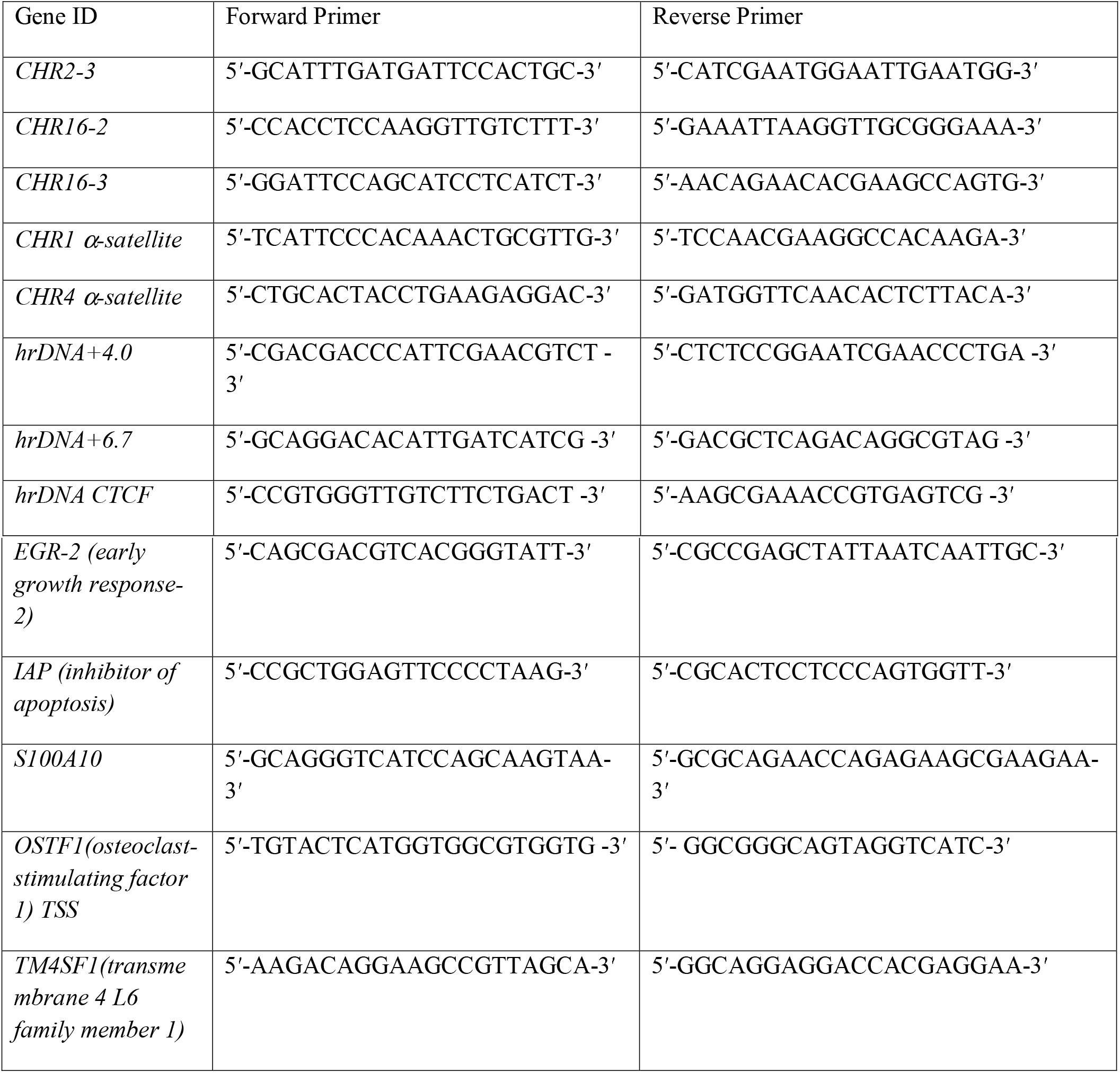

### Cell lysates and Western blot

Cells were washed with ice-cold PBS and collected by centrifugation. Whole cell lysates were prepared by using Triton X-100 lysis buffer (50 mM Tris-HCl, pH 7.4, 1% Triton X-100, 0.5% sodium deoxycholate, 0.1% SDS, 500 mM NaCl, 10 mM MgCl_2_, 10 mM sodium butyrate, and protease inhibitors). For Western blots, the samples were separated on 14% SDS PAGE, transferred to PVDF membrane and probed with corresponding antibodies.

### Cell fractionation, FLAG immuno-precipitation and mass spectrometry

Histone acetylation is crucial for its transportation, import into nuclei and deposit into chromatin. Therefore, it’s interesting to investigate how H3.3K56 mutations affect the dynamic changes of H3.3 associating proteins which plays critical roles in dominating H3.3 fate in the cell. To clarify this, H3.3WT, H3.3K56R and H3.3K56Q cells were collected, and cytosolic, nuclear extracts and chromatin binding fractions were extracted as previously described [49], and then FLAG immuno-precipitation was performed. To assess globally profile changes of H3.3 binding proteins in H3.3WT, H3.3K56R and H3.3K56Q cells, FLAG IP-purified H3.3 interacting proteins were subjected to 2D LC-MS/MS analysis and label free quantification as described [50]. The proteins were eluted from beads by boiling for 5 min, separated on SDS-PAGE followed by the MS-compatible Coomassie brilliant blue staining and in gel digestion. Briefly, the protein bands of interest were excised from the gel and cut into approximately 1mm^3^ slices. The gel slices were washed with 50 mM ammonium bicarbonate:acetonitrile (1:1) three times, SpeedVac-dried and digested by trypsin at 1:50 trypsin-to-protein mass ratio for overnight at 37 °C. The resultant peptides were extracted three times from the gel by vortexing in 10% formic acid:acetonitrile (1:1) for 15 min, and then dried by SpeedVac. The peptides were resuspended with 15 μL of 3% acetonitrile 2% formic acid before being subjected to nanoLC-MS/MS analysis.

#### LC-MS/MS analysis

MS data acquisition was performed by LC-MS/MS using a nanoLC.2D (Eksigent Technologies) coupled with a TripleTOF 5600+ System (AB SCIEX, Concord, ON). Peptides were directly loaded onto a reversed-phase Trap column (Chrom XP C18-CL-3μm 120A, Eksigent), separated with a linear gradient of 4–22% solvent B (0.1% FA in 98% ACN) for 50 min, 22–35% solvent B for 12 min and climbing to 80% solvent B in 4 min then held at 80% solvent B for the last 4 min, all at a constant flow rate of 300 nL/min on an Eksigent NanoLC 2D system. The resulting peptides were analyzed by Triple-TOF 5600+ mass spectrometer (AB Sciex). MS1 spectra were collected in the range 350–1,500 m/z for 250 ms. The 50 most intense precursors with charge state 2–5 were selected for fragmentation, and MS2 spectra were collected in the range 100–2,000 m/z for 100 ms; precursor ions were excluded from reselection for 15 s.

#### Database searching

The resulting MS data was processed using ProteinPilot (v2.3, AB Sciex). Tandem mass spectra were searched against UniProt_Human database (155,507 sequences) concatenated with reverse decoy database. Trypsin/P was specified as cleavage enzyme allowing up to 3 missing cleavages, 4 modifications per peptide and 5 charges. Mass error was set to 20 ppm for first search, 5 ppm for main search and 0.02 Da for fragment ions. Carbamidomethylation on Cys was specified as fixed modification and oxidation on Met, phosphorylation on Ser, Thr, Tyr and acetylation on protein N-terminal were specified as variable modifications. False discovery rate (FDR) thresholds for protein, peptide and modification site were specified at 1%. Minimum peptide length was set at 7. All the other parameters in ProteinPilot were set to default values. The site localization probability was set as > 0.75. All the other parameters in ProteinPilot were set to default values. In addition, ProteinProspector (v 5.20.0, UCSF) was used for search compare and protein quantification as described previously.

#### Label free quantification based on NSAF

Label free quantification of H3.3WT, H3.3K56Q or H3.3K56R associating proteins were performed as described. In brief, the NSAF (normalized spectral abundance factor) of a specific protein in each group (e.g. WT, Q and R) was used to represent their abundance, fold change of protein level in Q or R compared to WT was calculated by (NSAF in Q)/ (NSAF in WT), or (NSAF in R)/ (NSAF in WT), respectively. Proteins with spectra count >2 and fold change >1.5 or <0.66 were selected as significantly up- or down-regulated proteins and used for further Gene Oncology and KEGG pathway analysis.

#### Bioinformatics analysis

Heatmap (the R package “pheatmap”) for all quantified proteins were performed to determine the overall changes of H3.3WT, H3.3K56Q and H3.3K56R associating proteins in cytosolic fraction, nuclear extracts, or chromatin binding fraction. The differentially associating proteins were classified by Gene Ontology (GO) and Kyoto Encyclopedia of Genes and Genomes (KEGG) pathway annotation using DAVID online tools (http://david.abcc.ncifcrf.gov), and further demonstrated by heatmap. Protein-protein interaction network analysis was performed via STRING online tools (http://string-db.org), and manually reorganized.

## Results

### H3.3K56R mutant inhibits cell proliferation and causes cell death

To investigate the function of acetylation of histone variant H3.3 at lysine 56 (H3.3K56Ac), plasmids expressing H3.3K56Q or H3.3K56R mutations were constructed by site-directed mutagenesis, respectively, to mimic or antagonize H3.3K56 acetylation. DNA sequencing confirmed that the original code K (AAG) at position 56 was successively mutated to Q (CAG) or R (CGG), respectively (Fig 1A). FLAG-tagged H3.3WT (wild type), H3.3K56Q or H3.3K56R was then stably transfected into HeLa cells. The expression of FLAG-tagged H3.3K56WT, H3.3K56Q and H3.3K56R was determined by western blot analysis using antibodies against FLAG (Fig. 1B). Notably, while H3.3WT and H3.3K56Q stable cell lines grow relatively normally, H3.3K56R-transfected cells died gradually. The cell morphology of H3.3K56R cells was changed dramatically under microscope when compared with H3.3WT and H3.3K56Q stable cell lines (Fig. 1C). “Giant” cells with multi-nuclei were detected in H3.3K56R-transfected cells, suggesting that the formation of polyploidy may contribute to cell death induced by constant expression of H3.3K56R mutant (Fig. 1C).

**Fig 1.**
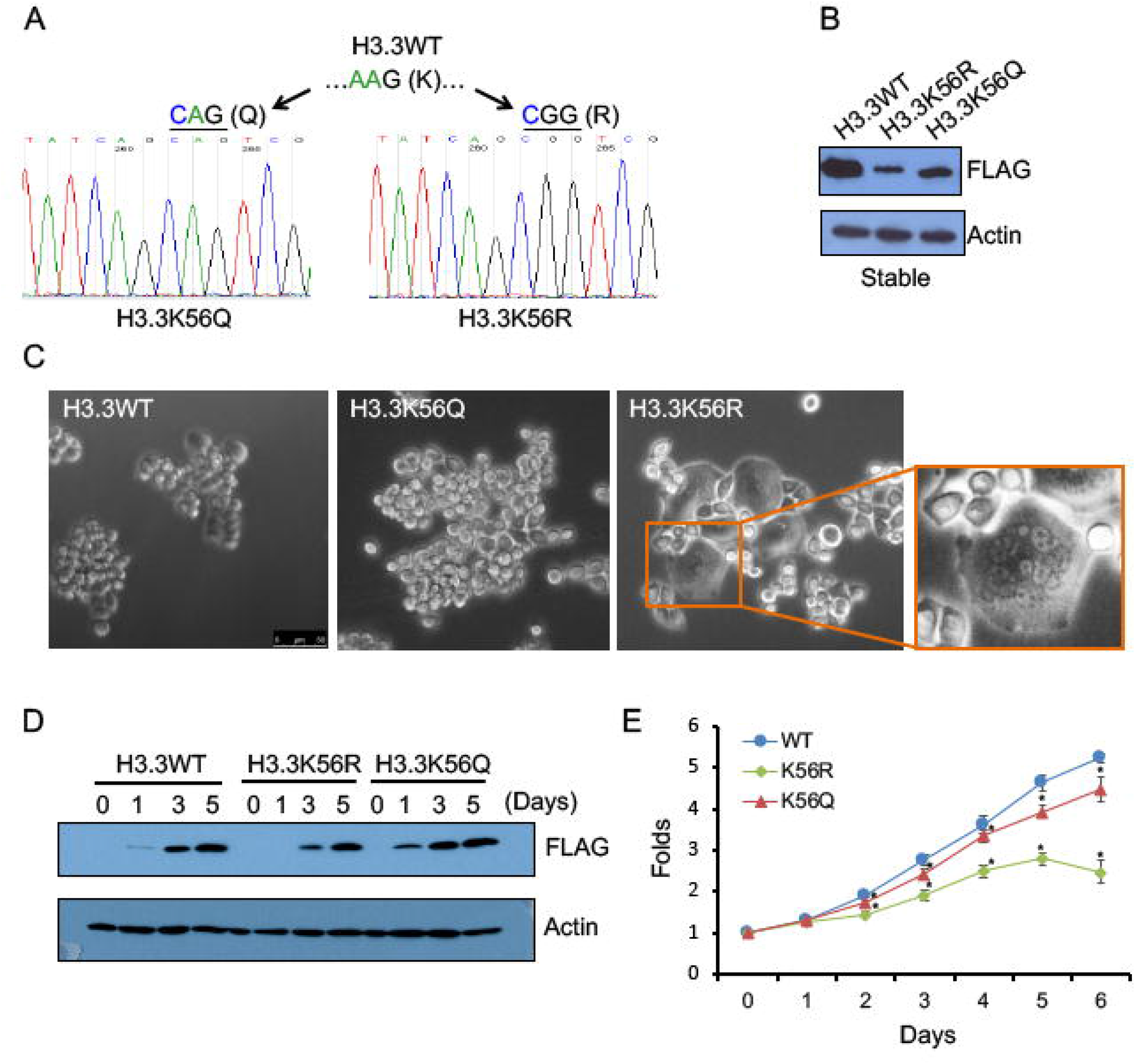
H3.3K56R mutant inhibits cell proliferation and causes cell death. (A) DNA sequencing results of H3.3K56 mutation to Q or R. Coding sequence AAG of H3.3K56 was substituted to CAG (Q) or CGG (R), respectively. (B) Expression analysis of H3.3WT, H3.3K56Q, or H3.3K56R in HeLa stable cell lines by western blot analysis using α-FLAG; α-Actin was used as loading control. (C) Effects of H3.3K56 mutation on morphology of HeLa cells. The morphology of H3.3WT and H3.3K56Q stable cells were ‘normal’, while H3.3K56R formed polyploidy cells. Notably, only H3.3WT and H3.3K56Q yielded final stable cell lines, H3.3K56R cells died gradually during the selection. (D) Inducible expression analysis of H3.3WT, H3.3K56Q, or H3.3K56R in UTA6 cells. FLAG-tagged H3.3WT, H3.3K56Q, or H3.3K56R were transfected into UTA6 cells with tet-off system to generate inducible stable cell lines. The expression of H3.3WT, H3.3K56Q, or H3.3K56R were induced by removal of tetracycline for 1, 3, and 5 days, and expression level was determined by western blot analysis using indicated antibodies. (E) UTA6-H3.3K56R mutant significantly decreases cell viability. UTA6 cells harboring inducible H3.3WT, H3.3K56Q, or H3.3K56R were induced expression for 0, 1, 2, 3, 4, 5 and 6 days, and subjected to viability assay using MTS method. The cell viability was normalized with day 0 control. The experiments were repeated at least three times. Means ± Standard Deviations (SDs); n=3; Student’s t-test, *:*p*<0.05.

Since constant expression of H3.3K56R mutant caused cell death, to study the function of H3.3K56Ac, the tet-off inducible cell lines were established that carry H3.3K56Q or H3.3K56R mutant (Fig. 1D). The plasmid containing wild-type or mutant K56 was transfected into UTA6 human osteosarcoma cells, which contain tetracycline-transactivator tTA. The stable cell lines were generated by the use of selection marker in the presence of tetracycline. The expression of H3.3WT, H3.3K56Q and H3.3K56R was induced by removing tetracycline from the culture medium and confirmed by western blot analysis using FLAG antibody (Fig. 1D). Next investigated were the effects of H3.3K56 mutations on cell proliferation using cell viability assays. As shown in Figure 1E, both K56R and K56Q mutants exhibited slower cell proliferation as compared with the control, albeit the proliferation of H3.3K56R mutant cells was more markedly inhibited than the K56Q mutant. For instance, on day 5 of the induction, the number of H3.3K56WT cells increased by 4.7-fold compared to day 0, whereas H3.3K56Q increased by 3.9-fold and H3.3K56R increased only by 2.7-fold (Fig. 1E). These results implicate a potential role for H3.3K56 acetylation in proper cell growth and cell division.

### polyploidy formation in H3.3K56R mutant cells

H3.1K56R mutant has been reported to cause S phase arrest and defects in cell cycle progression from S phase to M phase [37, 43]. To examine whether H3.3K56Ac plays similar roles as H3.1K56Ac in regulating cell cycle progression in mammalian cells, cell cycle analysis by flow cytometry following induction of H3.3K56 mutants in UTA6 cells for 3 days was performed. The cell cycle distribution was comparable between H3.3WT and H3.3K56Q cells, whereas H3.3K56R cells revealed reduced G1 population and increased polyploidy (Fig. 2B). Compared to H3.3K56WT control, the percentage of H3.3K56R cells with polyploidy was increased from 24% to 32%, while it remained unchanged in H3.3K56Q cells (Figure 2A). This is in line with the cell morphology changes observed in HeLa cells that were stably transfected with H3.3K56R mutant (Fig. 1C).

**Fig 2.**
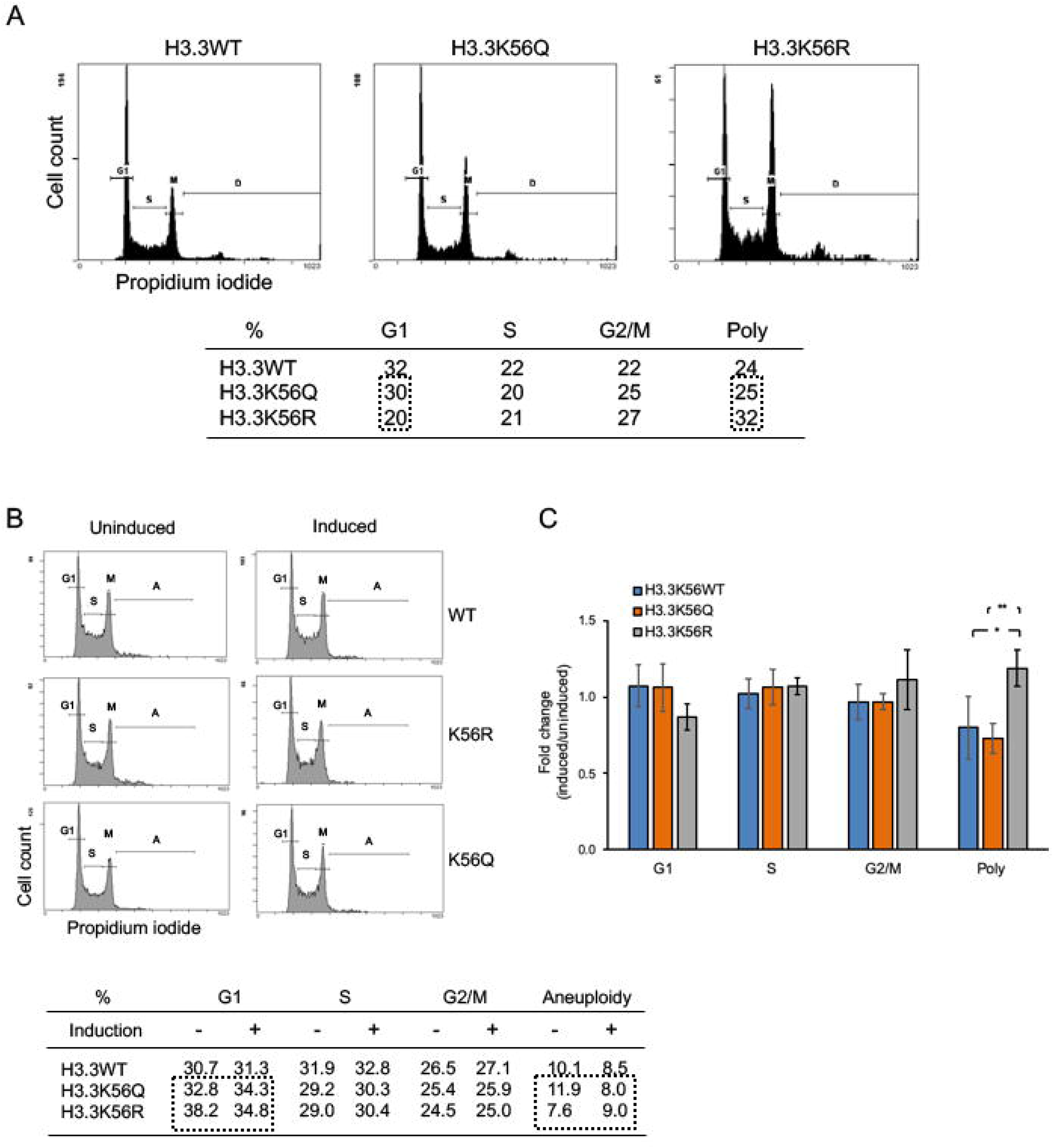
Polyploid formation in H3.3K56R mutant cells. (A) Cell cycle distribution analysis of HeLa cell lines stably expressing H3.3K56WT, H3.3K56Q, or H3.3K56R. (B) Cell cycle distribution analysis of UTA6 cells inducibly expressing H3.3WT, H3.3K56Q, or H3.3K56R. The expression was uninduced or induced for 3 days, and cells were collected for DNA content analysis by FACS. (C) Induced/uninduced fold changes of G1, S, G2/M or polyploid/aneuploidy population in UTA6 cells inducibly expressing H3.3WT, H3.3K56Q, or H3.3K56R. UTA6-H3.3K56R cells show significant higher induced/uninduced polyploid/aneuploidy population compared to both UTA6-H3.3WT and UTA6-H3.3K56Q. The data were presented as mean ±S.D. from the experiments performed in triplicate. *: *p*<0.05; **: *p*<0.01.

To rule out the possibility that observed polyploid cell population was simply resulting from overexpression of H3.3, regardless of its acetylation status at lysine 56, cell populations carrying aneuploid/polyploid before and after induction of H3.3K56WT, H3.3K56Q or H3.3K56R were compared. Before induction, ~8-12% of cells displayed aneuploidy/polyploidy in the H3.3K56WT, H3.3K56Q and H3.3K56R mutant cells (Fig. 2B). Following induction of the expression, the number of aneuploid/polyploid cells was increased in the H3.3K56R cells, while it was decreased in both H3.3K56WT and H3.3K56Q cells (Fig. 2B), suggesting that it is the lysine acetylation status at position 56 rather than overexpression of H3.3 itself important for the formation of aneuploidy/polyploidy. To determine if the changes are statistically significant, the ectopic expression of H3.3K56WT, H3.3K56Q and H3.3K56R mutants was induced for 5, 6, and 7 days and the fold changes (induced/uninduced) of cell populations at different cell cycle phases among different cell lines were compared. The aneuploid/polyploid cells were significantly increased in H3.3K56R cells after induction as compared to either H3.3K56WT or H3.3K56Q mutant cells (Fig. 2C). These results suggest that H3.3K56 acetylation is important for maintaining appropriate chromosome segregation and cell division.

### H3.3K56 acetylation regulates cell cycle progression through S phase and transition from S phase to G2/M phase

H3.1K56Ac mutants have been reported to cause S phase arrest and defects in cell cycle progression from S phase to M phase [37, 43]. While cell population in S phase was not changed by H3.3K56 mutants compared to H3.3K56WT (Fig. 3), it was not clear if these mutants have impact on progression through S phase in mammals. To study the role of H3.3K56Ac in cell cycle progression through S phase, H3.3WT, H3.3K56R, and H3.3K56Q cells were synchronized at the G1/S interface by double-thymidine block. After releasing the cells from the block for 2, 5, 8 and 12 hours, cell cycle analysis by flow cytometry was carried out. Two hours after the release, the percentages of cells in S phase were almost the same (62-65%) among the three cell lines. This made it possible to compare changes in cell population at different stages after 2 hours of release (Fig. 3A).

**Fig 3.**
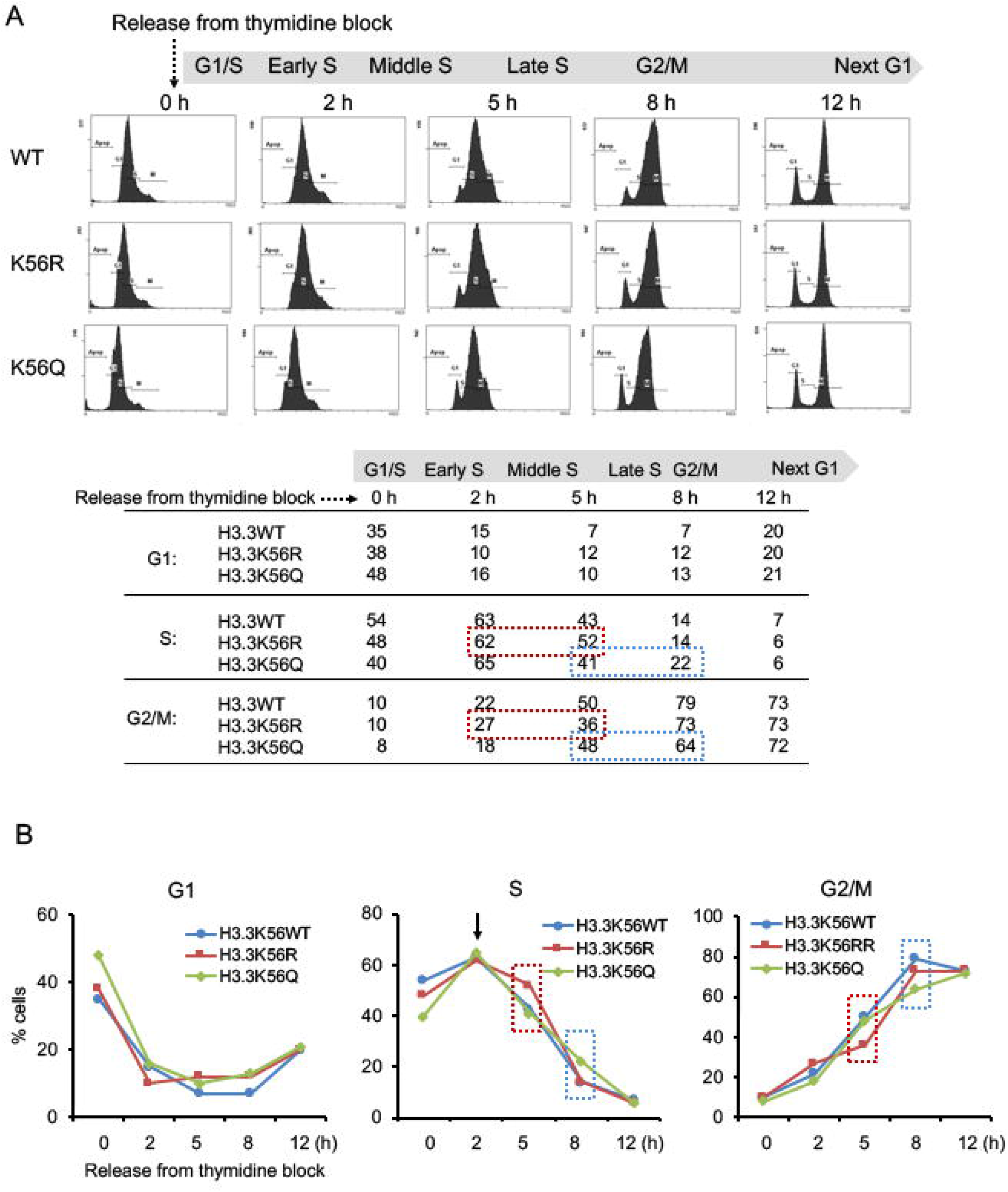
Abnormal cell cycle progression of H3.3K56R and H3.3K56Q mutant cells. (A) The H3.3K56WT/K56R/K56Q cells were induced expression, and synchronized at the G1/S interface by double-thymidine block respectively, then released for 2, 5, 8 and 12 h to monitor the cell cycle progression. The cells were collected, PI stained, and subject to FACS analysis to determine their cell cycle progression changes. (B) The percentage of G1, S and G2/M phase cells were presented as line graphs, showing the cell cycle differences over time.

Interestingly, delays in cell cycle progress were observed in both H3.3K56R and H3.3K56Q mutant cells, albeit at different time points. The percentage of H3.3K56R mutant cells in S phase was 52% at 5 hours compared to H3.3K56Q or H3.3K56WT cells which showed 41 or 43% cell population in S phase (Fig. 3A). Meanwhile, only 36% of H3.3K56R mutant cell had entered into M phase as compared with 48 or 50% of H3.3K56Q or H3.3K56WT cells in M phase at 5-hour release (Fig. 3A), indicating that the progression from middle to late S phase is compromised by H3.3K56R mutation. The line graphs show the differences in number of cells over time (Fig. 3B). A similar change was observed with H3.3K56Q cells 8 hours after the release. In this case, more H3.3K56K56Q cells stayed at S phase than both H3.3K56K56R and H3.3K56K56WT cells (22% vs. 14%), and less K56Q cells entered into M phase then other two type of cells (64% vs. 73~79%) (Fig. 3A), indicating a defect in transition from S to G2/M in H3.3K56Q cells. These results suggest that acetylation of H3.3K56 is required for the cell cycle progression from middle to late S phase, whereas H3.3K56 needs to be deacetylated to allow the transition from late S to G2/M phase.

### H3.3K56 mutant cell lines are more sensitive to DNA damage

It has been reported that acetylation status of H3K56 in yeast and H3.1K56 in mammals are critical for genomic stability and DNA damage response [37]. To examine the effect of H3.3K56 acetylation on genome stability, the expression of H3.3K56WT, H3.3K56Q or H3.3K56R was induced by removing tetracycline from the medium for 3 days, followed by immunostaining with γ-H2A.X, a sensitive marker for DNA damage. Unexpectedly, no obvious γ-H2A.X foci were observed in all three cell lines (not shown).

Next the cells were treated with the DNA damaging-reagents CPT and MMS, respectively, before and after the induction to test how these cells respond to DNA damage. Before the induction, the cell viability decreased dose-dependently by either CPT or MMS treatment, and no differences were observed among three different cell lines (Fig. 4A, B). After the induction, however, H3.3K56R and H3.3K56Q mutants showed significantly less cell viability compared to the H3.3K56WT control, indicating that both H3.3K56R and H3.3K56Q mutants are more sensitive to DNA damage than the control cells (Fig. 4A, B). Interestingly, H3.3K56R and H3.3K56Q mutants responded differently to CPT and MMS treatment. H3.3K56R mutant was more sensitive to CPT than H3.3K56Q, while H3.3K56Q showed higher sensitive to MMS than H3.3K56R mutant (Fig. 4), perhaps this is due to the distinct mechanisms of these two DNA-damaging reagents. These data suggest that H3.3K56 acetylation is required for proper DNA damage response.

**Fig 4.**
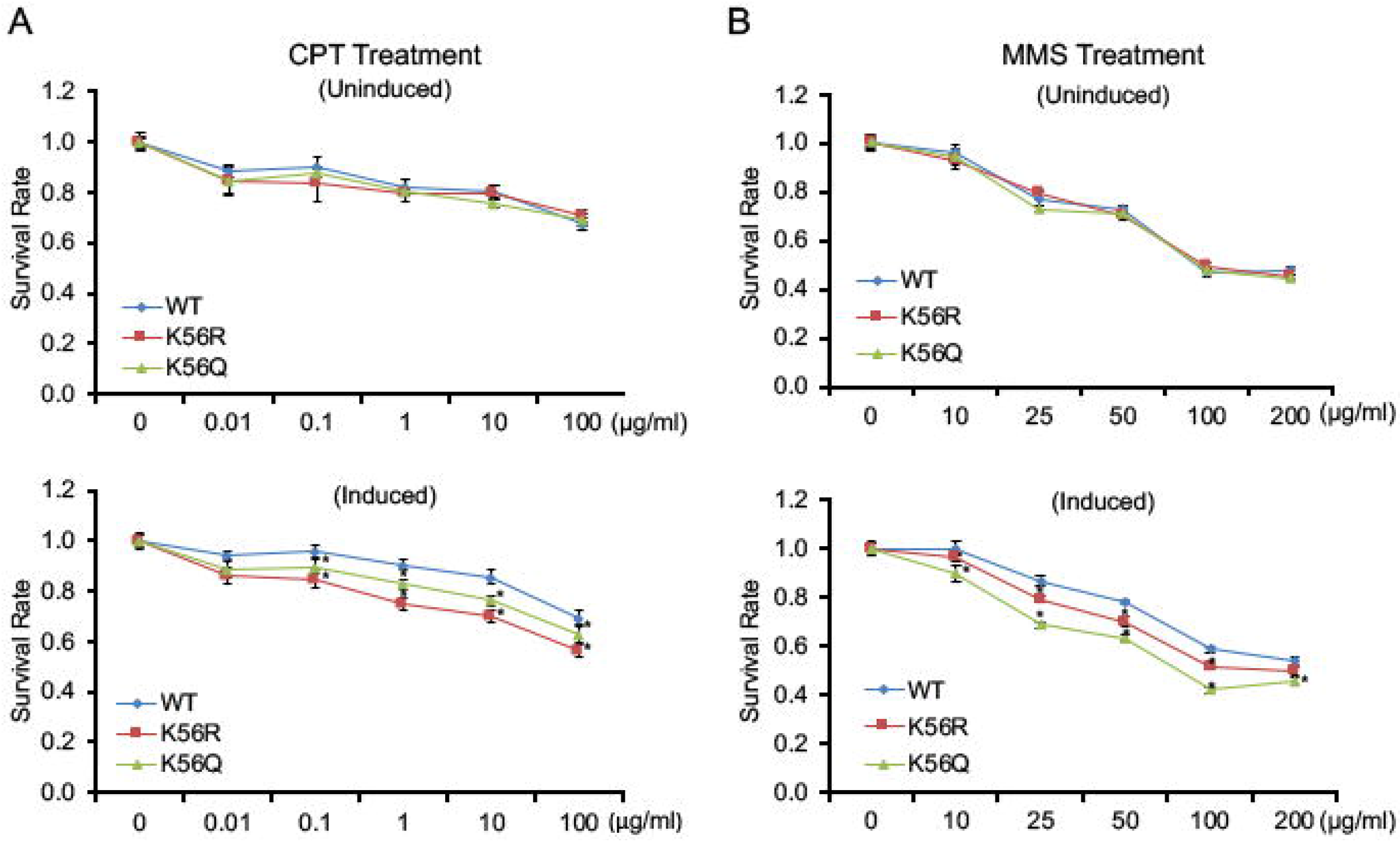
H3.3K56 mutant cells show differential sensitivity to DNA damage. UTA6 cells harboring inducible H3.3WT, H3.3K56Q, or H3.3K56R were induced expression for totally 3 days including treatment with CPT for 24 h (A) or MMS for 48h (B) at the indicated dosages. Cell viability was determined by the MTS assay. Both UTA6-H3.3K56Q and UTA6-H3.3K56R cells showed decreased viability compared to UTA6-H3.3WT. The cell viability was normalized with day 0 control. All the above experiments were repeated at least three times. Means ± SDs; n=3; Student’s t-test, *:*p*< 0.05.

### Distinct roles of H3.3K56 acetylation in H3.3 deposition

Lysine 56 acetylation promotes replication-dependent H3 assembly by enhancing the interaction between H3 and CAF-1 in yeast [43, 51]. To investigate whether K56 acetylation affects deposition of H3.3 into chromatin in mammals, ChIP-qPCR was performed to determine distribution of H3.3K56WT, H3.3K56Q and H3.3K56R mutants in genomic loci. The primers used covered a number of different genomic loci where H3.3 is localized, including enhancer/promoters, rDNA regions, pericentric and centric heterochromatin regions (Fig. 5A). ChIP-qPCR was carried out 1, 3, and 5 days after induction of ectopic expression of H3.3K56WT, and H3.3K56Q, or H3.3K56R mutants to monitor dynamic changes of their distribution. The H3.3K56Q mutant was enriched at some of pericentric (Chr2-3 and Chr16-2) and centric regions (Chr4) tested 3 days after the induction as compared with H3.3K56WT and the H3.3K56R mutant (Fig. 5C). This enrichment was not likely due to delayed release from the chromatin since (a) H3.3K56Q levels were much lower than those of H3.3K56WT and H3.3K56R at these sites after 1-day induction (Fig. 5B) and more importantly (b) there was no further increase of H3.3K56Q enrichment after 5-day induction (Fig. 5D). By contrast, the level of H3.3K56R mutant was remarkably decreased at Chr16-2 and Chr16-3 pericentric regions as well as Chr4 centric region (Fig. 5D). These results suggest that H3.3K56 acetylation is required for deposition of H3.3 at heterochromatin regions including some of pericentric and centric regions.

**Fig 5.**
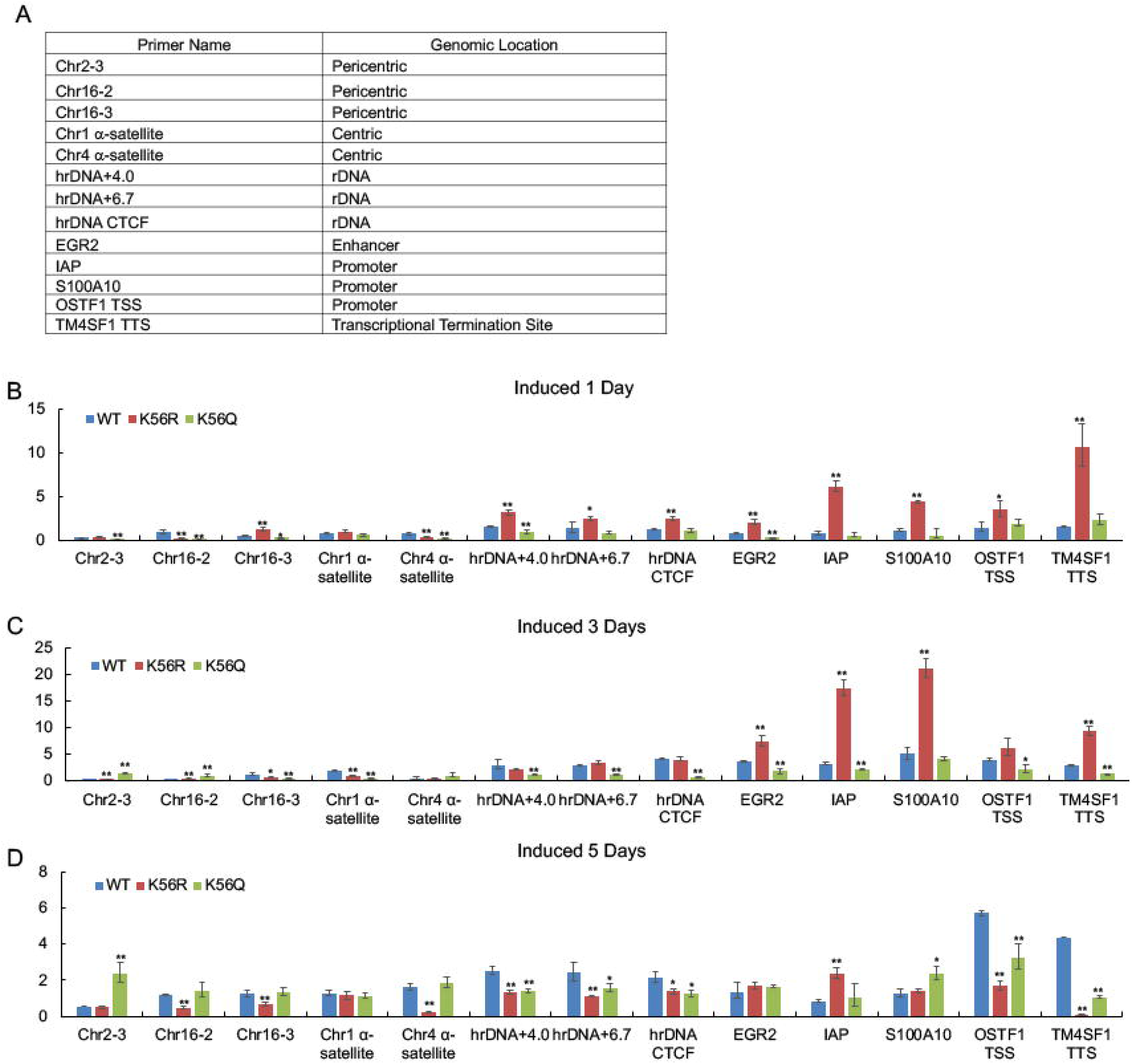
H3.3K56R mutants affect H3.3 assembly at distinct genomic loci. (A) Summary of tested primers and their corresponding genomic locations. (B-D) UTA6 cells harboring inducible H3.3WT, H3.3K56Q, or H3.3K56R were induced expression for 1, 3, and 5 days, mono- and dinucleosomes were isolated by Micrococcal Nuclease (MNase) digestion and sucrose gradient purification. Chromatin immunoprecipitation (ChIP) was performed using anti-FLAG M2 agarose beads, and the obtained ChIP-DNA was subject to qPCR using indicated primers representing different regions of genome, including pericentric, centric, rDNA, enhancer, promoter and transcriptional termination sites. Data are the Means ± SDs; n=3; Student’s t-test, *: *p*<0.05, **:*p*<0.01.

H3.3 was initially found localized at active rDNA regions, gene bodies of active genes and regulatory regions. To examine if K56 acetylation play roles in the deposition of H3.3 at these regions, ChIP-qPCR was performed with primers designed for several rDNA regions and around genes (Fig. 5A). Interestingly, the level of H3.3K56R was the highest and H3.3K56Q was the lowest among H3.3K56WT, H3.3K56Q and H3.3K56R mutants at most of these ‘active’ regions after 1-day induction (Fig. 5B), suggesting that K56 acetylation may inhibit incorporation of H3.3 into active rDNA and around active genes. This is supported by the ChIP-qPCR results obtained three day after the induction, which showed that while the amount of H3.3K56WT is increased compared to that from 1-day induction, reaching levels similar to the H3.3K56R mutant at rDNA regions in particular, the H3.3K56Q mutant levels did not changed at most of loci tested (Fig. 5C). The results obtained five days after the induction are more complicated. The level of both H3.3K56R and H3.3K56Q mutants were lower than that of H3.3K56WT in 5 out of 8 tested loci, including all three rDNA regions (Fig. 5D), indicating that H3.3K56 acetylation may inhibit both assembly and disassembly of H3.3 at these loci. Taken together, these data support the idea that H3.3K56Ac plays distinct roles in H3.3 assembly, either promoting or inhibiting the incorporation depending on the location and the timing.

### Distinct protein interaction network of H3.3K56R and H3.3K56Q mutants

To further understand mechanisms that underlie changes induced by H3.3K56 mutants, mass spectrometry was used to identify H3.3-associating proteins in different H3.3K56 mutant cell lines. The cytosolic fraction (Cyto), nuclear extracts (NE) and chromatin binding fraction (ChroB) were isolated from 3 day-induced H3.3K56WT, H3.3K56Q or H3.3K56R mutant cells, this was followed by FLAG immunoprecipitation to pull-down their associating proteins. After in-gel trypsin digestion, protein identification was carried out by nanoLC-MS/MS analysis (Fig. 6A). The heatmap showed distinct interacting-protein network for H3.3K56Q and H3.3K56R in each fraction (Fig. 6B-D). In general, H3.3K56Q- and H3.3K56R-associating proteins displayed opposite direction of the strength of the interaction in the cytosolic fraction and nuclear extracts in particular. A group of proteins in H3.3K56Q heatmap displayed in red (highest affinity), H3.3K56R in bright green (least affinity) and vice versa (Fig. 6B, C), suggesting a possibly opposite function of H3.3K56Q and H3.3K56R mutant.

**Fig 6.**
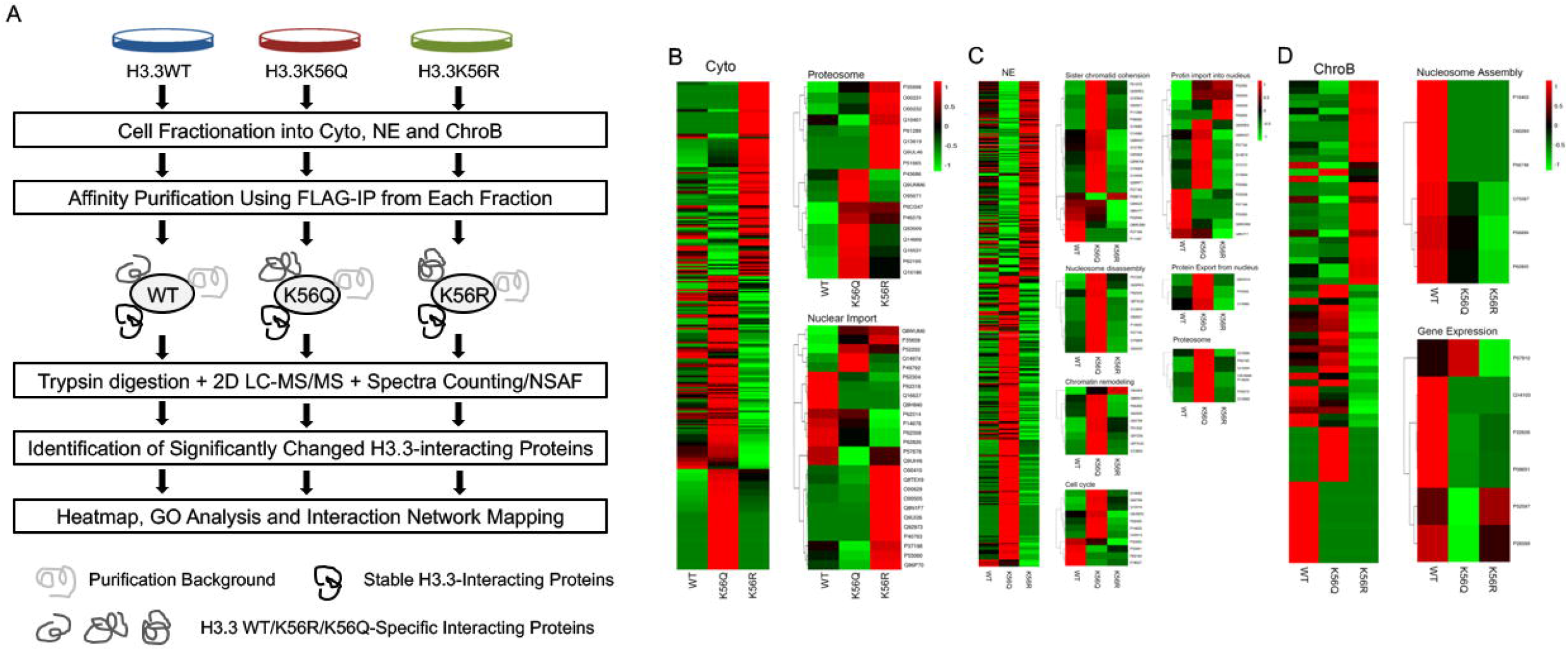
Distinct protein interaction network of H3.3K56R and H3.3K56Q mutants. (A) Schematic workflow of identifying significantly changed H3.3 interacting proteins of H3.3WT, H3.3K56Q and H3.3K56R by AP-MS. (B-D) UTA6 cells harboring inducible H3.3WT, H3.3K56Q, or H3.3K56R had induced expression for a total of 3 days, and then were fractionated into cytosolic, nuclear extract and chromatin binding fractions. Immune-precipitation with anti-FLAG M2 agarose beads was performed to purify H3.3WT, H3.3K56Q, or H3.3K56R interacting proteins in each fraction. The obtained protein complexes were subject to LC-MS/MS analysis and NSAF based label free quantification. Intensive bioinformatics analyses were performed for differentially binding proteins. Overall changes of H3.3WT, H3.3K56Q and H3.3K56R-associating proteins in cytosolic fraction (B), nuclear extracts (C), or chromatin binding fraction (D), respectively, as displayed by heatmap.

In cytoplasm, two major networks affected by the H3.3K mutation were “Proteasome” and “Nuclear Import”. H3.3K56Q and H3.3K56R mutations increased the binding of H3.3 to a distinct group of proteasome-associated proteins (Fig. 6B), indicating that both H3.3K56Q and H3.3K56R may be toxic to the cells and targeted by ubiquitin proteasome complexes. The changes in binding of the H3.3 mutants to proteins associated with nuclear import were more complicated. Notably, while H3.3K56R associated more strongly than H3.3K56WT with Importin 9 (Q96P70), Exportin 2 (P55060), Nuclear pore glycoprotein p62 (P37198), Importin 5 (O00629), and Importin 4 (Q8TEX9), whereas the binding of these proteins to H3.3K56Q was decreased or unchanged as compared with the H3.3WT (Fig. 6B). This suggests that H3.3K56 acetylation may inhibit the interaction between H3.3 and the proteins related to nuclear import. In the nuclear extract, the association between H3.3K56Q and proteins associated with sister chromatid cohesion, nucleosome disassembly, chromatin remodeling, cell cycle, protein import into nucleus, and protein export from nucleus, and proteasome, was greatly increased (Fig. 6C), while the association with H3.3K56R remained mostly decreased or unchanged. In the chromatin fraction, binding of H3.3 mutants to proteins responsible for gene expression and nucleosome assembly was inhibited in both H3.3K56Q and H3.3K56R cells with few exceptions (Fig. 6D). Taken together, these results suggest that H3.3K56Ac is critical for regulating H3.3-protein interactions, which may account for the observed phenotypes of H3.3K56 mutant cells.

The H3.3-protein interactions affected by both H3.3K56Q and H3.3K56R mutants were most apparent in the nuclear extract fractions (Fig. 6). To further characterize proteins and networks that are possibly regulated by H3.3K56 acetylation in the nuclear extract, a criteria for fold changes was determined, i.e., K56Q VS. WT > 1.5 and K56R VS. WT < 0.67, or K56Q VS. WT < 0.67 and K56R VS. WT > 1.5 to identify significantly dysregulated H3.3K56Q- and H3.3K56R-binding proteins that meet the criteria. Proteins identified based on a change in an opposite direction by K56Q and K56R are more likely those whose interaction with H3.3 was regulated by K56 acetylation. Then Gene Oncology and protein-interaction network analyses were performed. In a set of proteins whose association with H3.3 was up-regulated by H3.3K56Q but down-regulated by H3.3K56R, the most enriched biological processes included sister chromatid cohesion, mitotic nuclear envelope disassembly, protein import into nucleus, and mitotic nuclear division (Figs. 7A, B and Table 1). On the other hand, rDNA processing and nucleosome assembly were among the most enriched biological processes in H3.3K56Q down-regulated but H3.3K56R up-regulated proteins (Figs. 7C, D and Table 1). These results indicate that acetylation status of H3.3K56 appears to be critical for proper chromosome segregation during cell division, which may explain the failure of chromosome segregation and formation of polyploid/aneuploid in the H3.3K56R mutant cells.

**Fig 7.**
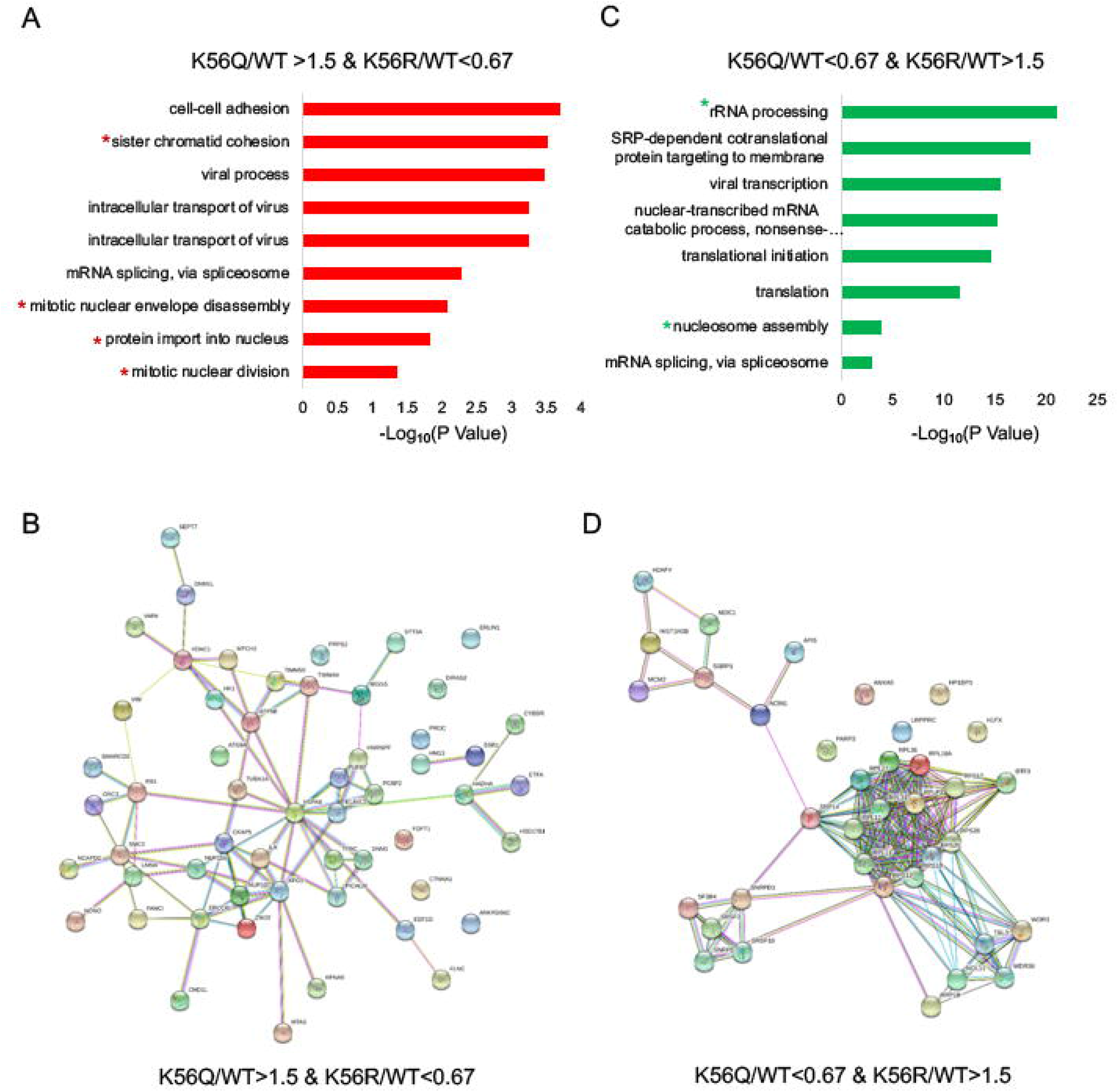
Significantly changed H3.3K56Q and H3.3K56R-associating proteins in nuclear extracts. Biological processes enriched in GO analysis for differential associating proteins were up-regulated in H3.3K56Q but down-regulated in H3.3K56R (A) or down-regulated in H3.3K56Q but up-regulated in H3.3K56R (C), respectively, in nuclear extract. (B and D). Protein interaction network of proteins in A or C, respectively.

**Table 1.**
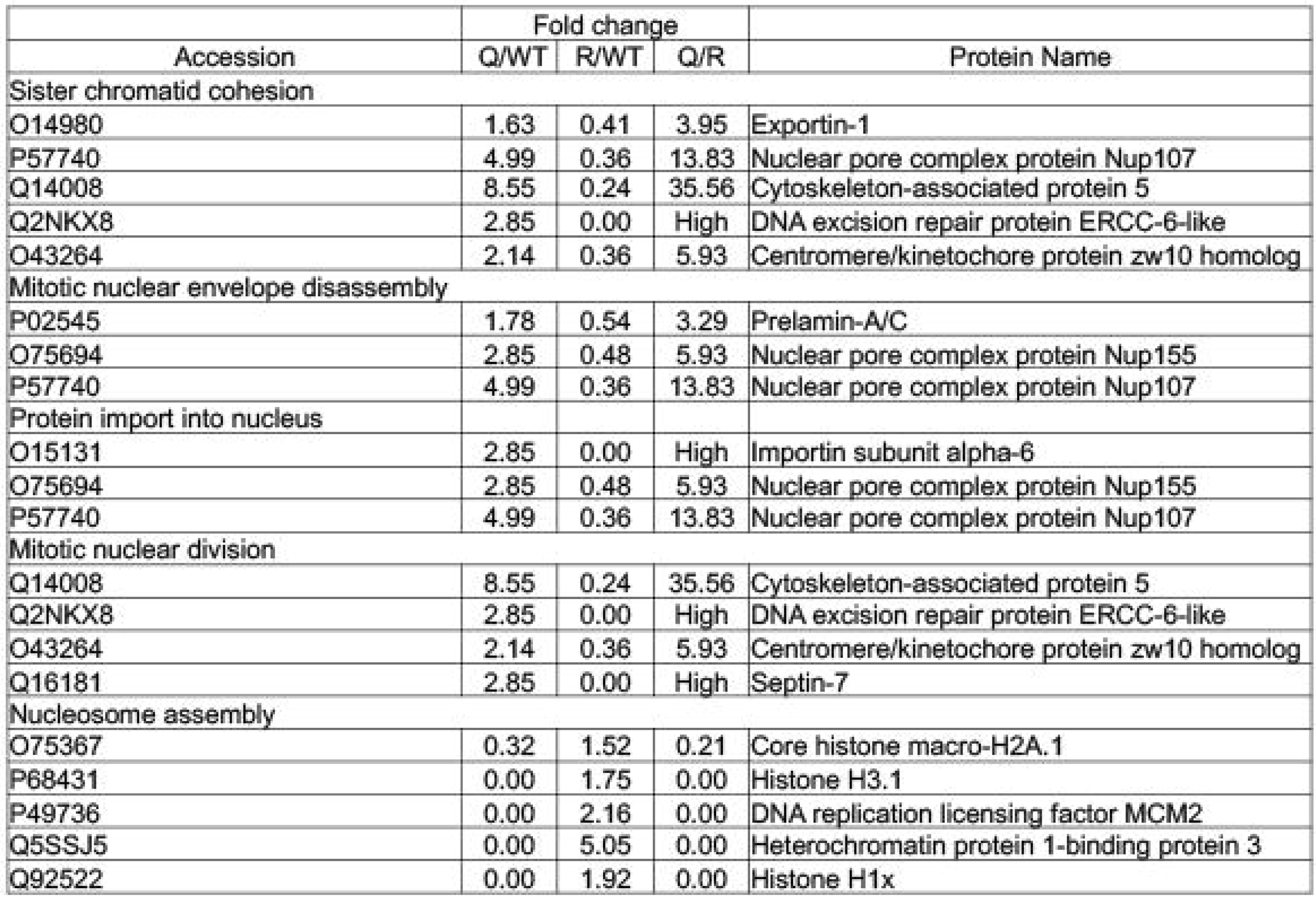
Selected significantly changed biological processes enriched from H3.3K56Q and H3.3K56R-associating proteins in nuclear extracts. Up-regulated biological processes: sister chromatin cohesion, mitotic nuclear envelope disassembly, protein import into nucleus and mitotic nuclear division; Down-regulated biological processes: nucleosome assembly. The corresponding protein name, accession number and fold changes between Q/WT, R/WT and Q/R were also shown.

## Discussion

### Potential role of H3.3K56 acetylation in cell division and chromosome segregation

Since its first finding in yeast, H3K56Ac has been identified to be conserved in various species including *Drosophila* and mammals. Studies in mammals demonstrated that H3.1K56ac plays important roles in maintaining genomic stability, DNA replication, cell cycle progression and DNA repair [30, 35, 45]. While H3.1K56Ac has been relatively well studied, the function of H3.3K56Ac has remained largely unknown.

Pal *et al*. reported that the commercial antibodies against H3K56 acetylation are nonspecific in human cells [52], which made it challenging to study this modification. It would be more challenging to develop antibodies specific for H3.3K56 acetylation given a few amino acid differences between H3.3 and canonical H3. Thus, to study the function of H3.3K56 acetylation, H3.3K56Q and H3.3K56R mutants were made to mimic or prevent acetylation at position 56, respectively. The first distinct change observed was large multinucleated cells which were consistently expressed the H3.3K56R mutantion (Fig. 1C). By contrast, H3.3K56Q and H3.3K56WT cells were morphologically normal. Multinucleation can occur by fusion or failure of cytokinesis. In a study using murine fibroblasts, Holt *et al*. found that multinucleation is produced by fusion only in the presence of macrophages [53]. In the absence of co-cultured macrophages, these cells multinucleate not by fusion, but from senescent cells that no longer undergo cytokinesis. Therefore, the observed “giant” multinucleated cells stably expressing the H3.3K56R mutant may represent senescent cells undergoing mitosis without cytokinesis. This is consistent with the observation that the cells consistently expressing H3.3K56R eventually died during long-term culture. Thus, these results indicate that H3.3K56 acetylation might be required for proper cytokinesis.

Since the cells that stably express H3.3K56R mutant died gradually, to precisely investigate the function of H3.3K56Ac, a tet-off system was used to generate the cell lines in which the expression of H3.3K56WT, H3.3K56Q or H3.3K56R was inducible. Notably, although FACS analysis showed that cell cycle progression of both H3.3K56WT and H3.3K56Q cells was similar, H3.3K56R cells displayed reduction of G1 phase duration and increase in polyploidy formation (Fig. 2A). Moreover, while the polyploidy/aneuploidy population was decreased following induction of both H3.3K56WT and H3.3K56Q mutant, it was increased upon induction of H3.3K56R, the difference was statistically significant (Fig. 2C). This supported the idea that H3.3K56 acetylation is important for cell division/chromosome segregation.

This conclusion is also evidenced by the results showing the changes in H3.3 deposition and H3.3-protein interactions by the H3.3K56 mutants. Deposition of H3.3K56R was compromised at several of centric and pericentric regions tested compared with H3.3WT or H3.3K56Q, in particular five days after the induction (Fig. 5). It is known that during S phase, H3.3 is deposited at centric domain as a ‘placeholder’, which is replaced later with CENP-A in G1 phase [54]. Centromeres consist of the centric heterochromatin, which ensures kinetochore formation, and the pericentric heterochromatin, which is responsible for the chromatid cohesion [55]. Interestingly, the mass spectrometry data demonstrate that the proteins involved in the processes of sister chromatid cohesion, mitotic nuclear envelope disassembly and mitotic nuclear division, including centromere/kinetochore protein zw10 homolog, cytoskeleton-associated protein 5 (CKAP5), and septin-7 were among the proteins whose association with H3.3 was up-regulated by the K56Q substitution but down-regulated by the K56R substitution (Figs. 7A, B and Table 1). Centromere/kinetochore protein zw10 homolog is an essential component of the mitotic checkpoint, preventing cells from prematurely exiting mitosis [56]. CKAP5 plays an essential role in centrosome microtubule assembly[57]. Moreover, septin-7 is required for normal progress through mitosis and involved in cytokinesis [58]. Together, these results all point to a critical role for H3.3K56 acetylation in chromosome segregation and cytokinesis during cell mitosis. Disruption of acetylation at lysine 56 could lead to chromosome mis-segregation and subsequent polyploidy/aneuploidy.

### H3.3K56 acetylation and H3.3 assembly

H3K56Ac has been reported to promote efficient chromatin assembly during DNA replication in part by enhancing the binding of nucleosome assembly factors for newly synthesized histone H3 [31, 51]. H3K56Ac also promotes efficient release of newly synthesized histones from histone chaperones, such as Asf1, by facilitating transient ubiquitination of histone H3, thereby increasing the availability of free histones for downstream chaperones [59, 60]. In DNA repair, H3K56Ac on free histones is required for chromatin assembly after repair, possibly by promoting H3 recruitment to the repaired DNA by increasing its binding affinity to repair specific histone chaperones [41]. The mass spectrometry analysis coupled with label free quantification implies that H3.3K56Ac may also play an important role in regulating H3.3-binding partners responsible for nucleosome assembly. For instance, in nuclear extract, H3.3K56Q increased association of H3.3 to proteins associated with nucleosome disassembly and chromatin remodeling, along with others, while H3.3K56R increased interaction between H3.3 and proteins responsible nucleosome assembly (Figs. 6 and 7).

Marked changes in H3.3K56Q and H3.3K56R binding proteins related to nucleosome assembly, suggest potential effect of K56 acetylation on the deposition of H3.3 into the chromatin. In general, H3.3K56R showed much higher deposition into all ‘active’ sites tested, especially one day after the induction, including the rDNA regions and the enhancer/promoters (Fig. 5B). However, deposition of H3.3K56Q into most of these regions was significantly inhibited, especially three days after the induction (Fig. 5C). The majority of chromatin assembly/disassembly that occurs within one day induction could be considered replication-independent. Therefore, these results indicate that K56 acetylation may have negative impact on replication-independent assembly of H3.3. In contrast, H3.3K56Q increased its incorporation into heterochromatin regions tested three days after the induction, while deposition of H3.3K56R mutant into these regions was significantly inhibited, especially five days after induction (Fig. 5C, D). These data imply that H3.3K56 acetylation is required for assembly of H3.3 into certain heterochromatin regions. Together, these results suggest that H3.3K56Ac may play a critical role in switching between replication-dependent and replication-independent assembly of H3.3 at distinct genome loci.

### H3.3K56 acetylation in cell cycle progress and DNA damage response

Cell proliferation was inhibited by both H3.3K56Q and H3.3K56R mutations, with H3.3K56R showing a more significant inhibition (Fig. 1E). Interestingly, cell synchronization and release results demonstrated that H3.3K56Ac was not required in the early stage of G1/S to S transition, but was critical for the cell cycle progression from middle S to late S phase. Moreover, it seemed that H3.3K56Ac had to be deacetylated to allow the transition from late S into M phase (Fig. 3). These results are generally aligned with the pattern of H3K56 acetylation during S phase observed in the yeast and for canonical H3.1.

In yeast, accumulating evidence indicates that acetylation-deacetylation cycle of H3K56 is critical for efficient cellular responses to DNA damage as demonstrated by the dynamic switch of acetylation and deacetylation of H3K56 promoting cell survival in response to spontaneous or genotoxic agent-induced DNA lesions [36, 61]. It has been reported that HeLa cells expressing H3.1K56R and H3.1K56Q mutants were more sensitive to DNA damage caused by CPT, HU and MMS [37]. However, in this study only H3.3K56R and H3.3K56Q mutant cells showed higher sensitivity to CPT or MMS induced DNA damage, while H3.1K56R and H3.1K56Q mutant cells remained unaffected (Fig. 4 and data not shown). This suggests that H3.3K56Ac might play more critical role in DNA damage response than H3.1K56Ac. Unexpectedly, when comparing H3.3K56R to H3.3K56Q, H3.3K56R mutant cells were more sensitive to MMS, while H3.3K56Q cells were more sensitive to CPT, supporting that MMS and CPT might cause DNA damage by different mechanisms [62, 63]. Given that HeLa stable cell lines expressing H3.1WT, H3.1K56Q or H3.1K56R are all viable and grow ‘normally’ [37], these results suggest that H3.3K56Ac might possess a more essential function then H3.1K56Ac.

In summary, it was found that H3.3K56 acetylation may play important roles in many aspects of cellular processes including nucleosome assembly, normal cell growth, cell cycle progress, and DNA damage response in mammals. In particular, it was demonstrated that H3.3K56 acetylation appears to be critical for proper chromosome segregation and cell division. Through proteasome analysis, a spectrum of H3.3-associating proteins was identified whose interactions with H3.3 were significantly changed by abnormal acetylation status of H3.3K56. This provided a potential mechanistic insight into the function of H3.3K56 acetylation. Future study is needed to identify which H3.3-associating proteins are indispensable for H3.3K56 function. Also, study could explore the mechanisms that underlie potential function of K56 acetylation in determining replication-dependent or -independent assembly of H3.3.

## Funding supports

This work was supported by NYU Grossman School of Medicine Start-up package to C. J. and by the National Natural Science Foundation of China (Grant No. 31500664, 31770838) and Natural Science Foundation of Jiangsu Province (Grant No. BK20171338) to Lei Fang.

## Acknowledgements

We thank “Translational Medicine Core Facilities of Medical School of Nanjing University” for the use of mass spectrometry and bioinformatics analysis.

## Author Contributions

CJ and LF conceived and designed the experiments. LF, DC, JZ, and HL performed the experiments and analyzed the data. CJ, LF, DC, and BB wrote the paper.

